# C-terminally encoded peptides promoting root symbiotic nodulation in legume plants also promote the root arbuscular mycorrhizal symbiotic interaction

**DOI:** 10.1101/2024.07.17.603861

**Authors:** Léa Pedinotti, Juliette Teyssendier de la Serve, Thibault Roudaire, Hélène San Clemente, Marielle Aguilar, Wouter Kohlen, Florian Frugier, Nicolas Frei dit Frey

## Abstract

C-terminally encoded peptides (CEPs) are small secreted signalling peptides that promote in legumes the root nitrogen-fixing nodulation symbiosis depending on soil mineral nitrogen availability^1^. In Medicago truncatula, their action is mediated by the Leucine-rich repeat receptor-like protein kinase COMPACT ROOT ARCHITECTURE 2 (CRA2)^2–4^. As most land plants, under inorganic phosphate (Pi) limitation, M. truncatula establishes another root endosymbiotic interaction with arbuscular fungi, the arbuscular mycorrhizal symbiosis (AMS). Because this interaction is beneficial for the plant but has a high energetic cost, it is tightly controlled by host plants to limit AMS infections mainly depending on Pi availability^5^. We show in this study that the expression of a subset of CEP encoding genes is enhanced in the low Pi conditions that favour AMS colonization, and that overexpression of the low Pi-induced MtCEP1 gene, previously shown to promote the nitrogen-fixing root nodulation symbiosis, enhances the AMS colonization from the initial entry point of the fungi. Conversely, a loss-of-function mutation of the CRA2 receptor required for mediating CEP peptides action^2^ decreases the AMS interaction capacity from the same initial fungal entry stage. Transcriptomic analyses revealed that the cra2 mutant is negatively affected in the regulation of key Pi transport and response genes as well as in the biosynthesis of strigolactone (SL) hormones that are required for establishing the AMS interaction. Accordingly, SL contents were drastically decreased in cra2 mutant roots. Overall, we showed that the CEP/CRA2 pathway promotes, in legume plants, both root nodulation and AMS depending on soil mineral nutrients availability.

**In brief:** The establishment and maintenance of root nodule and arbuscular mycorrhizal symbioses (AMS) in legume plants is tightly regulated respectively by nitrogen and phosphate availability. Pedinotti and Teyssendier de la Serve et al. show that the CEP/CRA2 pathway, previously known to promote root competence to nodulate under low nitrogen conditions, also enhances AMS establishment by controlling the expression of Pi homeostasis and strigolactone (SL) hormone biosynthesis genes, as well as SL accumulation.

**Highlights:**

- A subset of M. truncatula CEP genes is induced by AMS-promoting low phosphate conditions.
- The CEP1 peptides positively regulate AMS establishment, while the cra2 CEP receptor loss of function mutants display a reduced AMS colonization.
- The CEP/CRA2-dependent regulation of AMS acts through downstream signals distinct from the ones involved in root nodule symbiosis regulation. The CEP/CRA2 pathway regulates AMS establishment by controlling Pi homeostasis and strigolactone (SL) hormone biosynthesis genes, as well as SL accumulation.

## Results and Discussion

### Low phosphate availability induces the expression of a subset of CEP genes

To determine whether CEP peptides can act in AMS, we analysed the regulation of all M. truncatula Class I CEP genes (Figure 1 and S1) using RT-qPCR in contrasting inorganic phosphate (Pi) availability conditions (low Pi = 7.5 µM; high Pi = 2 mM) or in arbuscular mycorrhizal fungi (AMF) colonized roots. None of the CEP genes were significantly regulated upon AMF colonization (Figure 1A and S2A), despite the strong induction of the PT4 mycorrhization marker (Figure S2B^6^), validating the efficient mycorrhization. In contrast, the expression of MtCEP1, MtCEP2, MtCEP4, MtCEP5, MtCEP6, MtCEP7 and MtCEP9 was significantly higher in the low Pi condition. Mt4, a Pi starvation inducible gene^7^, was used as a marker to validate that the Pi conditions were indeed contrasted (Figure S2C^8^), as also attested by the overall plant growth phenotype (Figure S3). These results indicate that the expression of a subset of CEP peptides encoding genes is promoted in low Pi conditions that are required for AMS establishment, suggesting a potential regulatory role of these signalling peptides in this symbiotic interaction. Interestingly, a role for Class I CEP peptides was previously identified for another legume root endosymbiotic interaction, the nitrogen-fixing root nodule symbiosis. The regulation of these genes by Pi availability, but not by the AMS interaction by itself^1,2^, echoes the current model of the CEP-mediated regulation of nodulation by nitrogen availability^4,9^. Overall, this suggests that CEP peptides may promote not only symbiotic root nodulation but also AMS.

**Figure 1.**
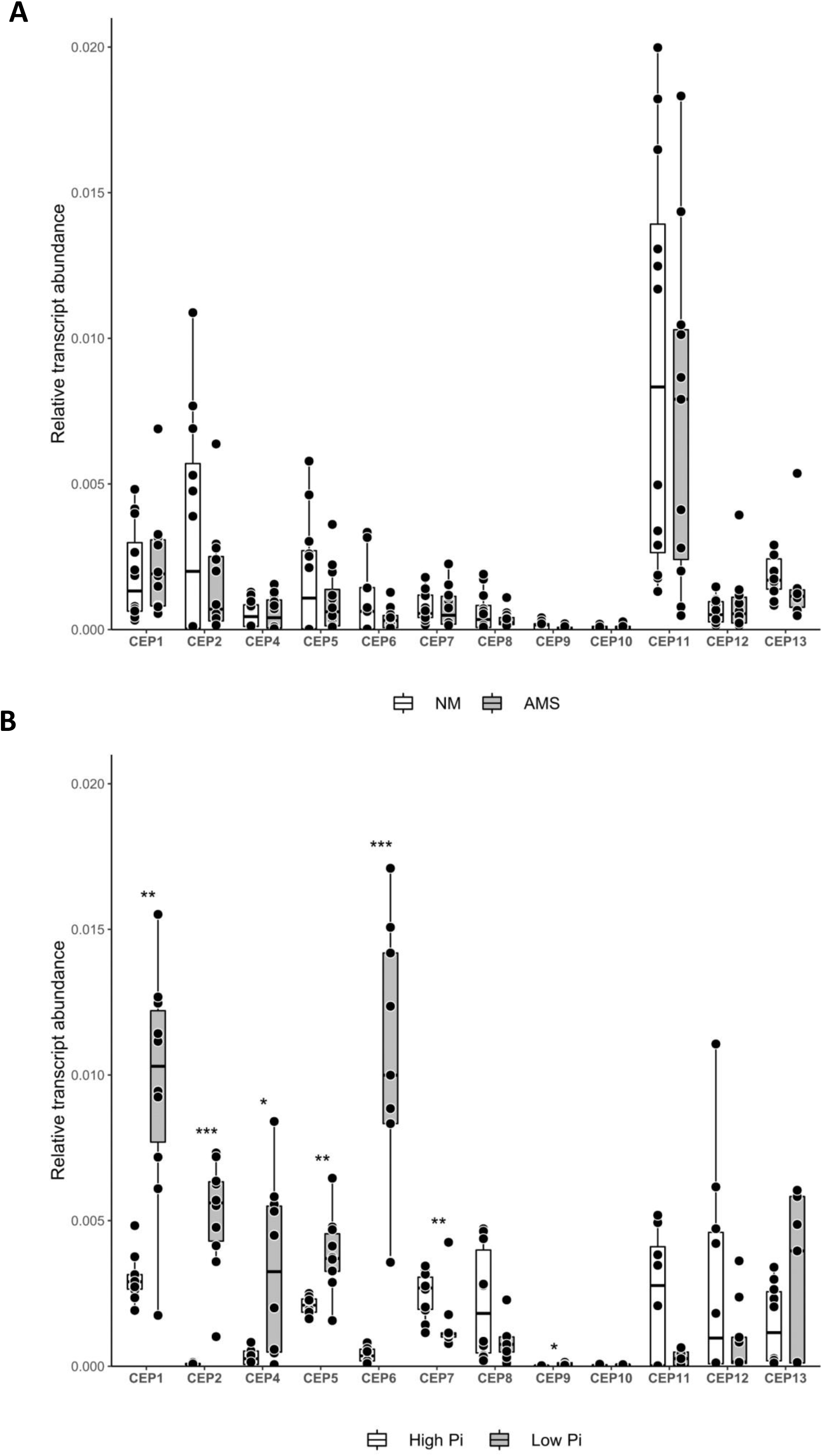
A subset of MtCEP genes is upregulated in low Pi conditions. **A.** RT-qPCR analysis of class I MtCEP gene expression with Arbuscular Mycorrhizal Symbiotic (AMS) fungus inoculation or without (NM, Non-Mycorrhized). A Mann-Whitney test performed between the two conditions (AMS and NM) revealed no significant difference for each gene (P < 0.05). **B.** RT-qPCR analysis of class I MtCEP gene expression in low and high Pi conditions. A Mann-Whitney test was performed between the two conditions (low and high Pi): *, P < 0.05; **, P < 0.01; ***, P < 0.001. Data from two independent biological experiments are shown (n=5 pools of 3 plants per experiment). Boxes represent the middle 50% of the values, the median is represented by a horizontal line, and upper and lower quartiles by vertical lines. Expression values were normalized to the MtACT11 gene.

### Overexpression of a CEP peptide enhances AMS root colonization whereas mutation of the CRA2 receptor conversely reduces AMS

With the aim of further comparing the role of CEP peptides in both root endosymbioses, we selected MtCEP1 among CEP genes responding to Pi availability, first because it was already studied in the symbiotic nodulation context using a 35S:MtCEP1 overexpression approach^1^; and second because MtCEP1 is one of the most expressed CEP gene in the low Pi condition (Figure 1B). Roots overexpressing MtCEP1 showed a significantly increased density of the different mycorrhization events quantified, from hyphal epidermal entry sites to hyphae root colonisation and arbuscule formation, despite root apices number were reduced, when compared to empty vector-transformed roots (Figure 2A). Overall, this indicates that, in addition to its known role as a positive regulator of nodulation^1,2^ and as a negative regulator of lateral root development^1^, MtCEP1 peptides also promote AMS. We additionally tested the mycorrhizal phenotype of CEP2 overexpressing roots, a CEP peptide encoding gene also induced by low Pi with a high fold change but reaching an overall expression level lower than MtCEP1 (Figure 1B). Similarly as for CEP1 overexpression, an increased number of infection points was observed (Figure S4A), which however did not translate into an increased overall colonization of roots by the fungi. This suggests that the different CEP genes accumulating in low Pi conditions might have, beyond the regulation of the initial fungi entry points in roots, differential functions on later mycorrhizal stages. As CEP peptides act through the CRA2 receptor in M. truncatula^2–4^, we additionally analysed the mycorrhization phenotype of cra2 mutants. In both cra2-11 (hereafter called cra2) and cra2-1 alleles, respectively generated in the Jemalong A17 or the R108 genotypes, the mycorrhizal colonization was similarly reduced (Figure S4B). Moreover, a detailed phenotyping of cra2-11 mutants revealed that epidermal entry sites, infection points, hyphae, and arbuscule rates were all reduced compared to WT plants (Figure 2B), while the lateral root number was increased in the mutant, as expected^10^. Careful observation of arbuscule using confocal microscopy however did not reveal major differences between cra2 mutants and the WT (Figure S5). This suggests that while cra2 mutants display less colonization events, once the mutant hosts the fungus fur nutrient exchanges, it is able to achieve arbuscule development similarly as in the WT. When normalizing the AMS colonization by the number of lateral roots, the mycorrhization phenotype of the cra2 mutant was even increased compared to WT plants (Figure S6). The increased number of young lateral roots, and thus of susceptible regions to AMS^11^, would be expected to increase its AMS colonization rate. In contrast, the opposite result is observed, indicating even more convincingly than without normalization that the cra2 mutant has a very low competency to interact with the AMS fungi. Of note, the same occurs for symbiotic nodulation: despite having more lateral roots containing young regions above root tips that are the most responsive to rhizobia nitrogen-fixing bacteria, the cra2 mutant forms nearly no symbiotic nodule^10^. As the different stages of AMS were impacted, either by the MtCEP1 overexpression or by the CRA2 loss of function mutation, this suggests that the CEP peptide / CRA2 receptor pathway is required to promote the establishment of the AMS from the earliest infection stage, similarly as previously highlighted for the nitrogen-fixing symbiotic nodule initiation^3,9^. This thus represents another example of the likely evolutionary recruitment of a more ancient regulatory pathway promoting AMS for the regulation of the more recently evolved nitrogen-fixing root nodule endosymbiotic interaction^12^. Such recruitment may have also involved a potential rewiring of CEP gene regulation depending on different environmental inputs as previously documented for the NIN/NLP (Nodule Inception / NIN-Like Protein) regulatory genes in the context of symbiotic nodulation^13,14^. As CEP genes are both induced by low nitrogen and low Pi environments, they might coordinate the root competency for both symbioses, depending on the combination of nutrient availabilities in the soil.

**Figure 2.**
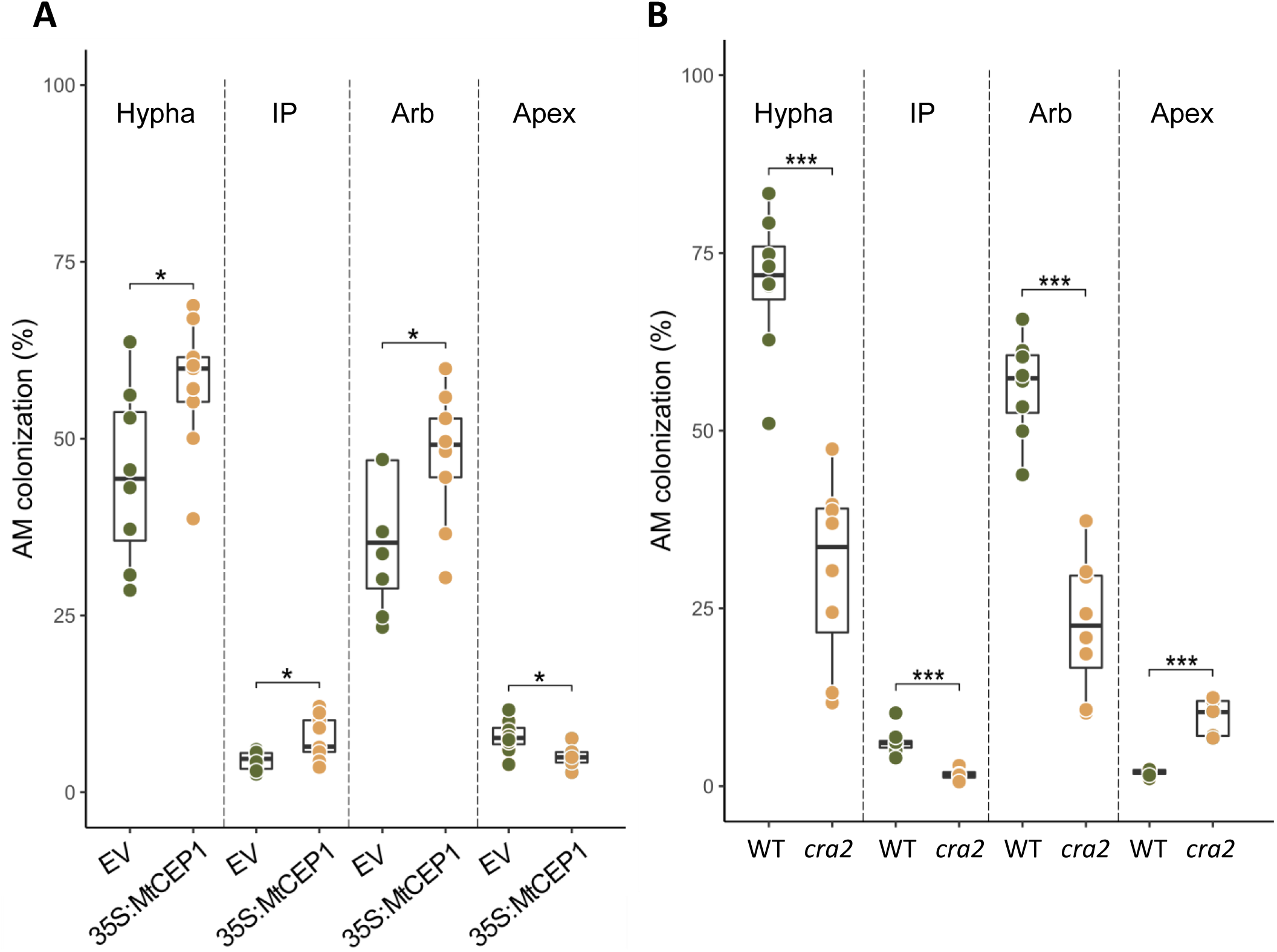
Mycorrhization phenotypes of roots overexpressing MtCEP1 and of the cra2 mutant. **A.** Mycorrhizal phenotype of MtCEP1 overexpressing (35S:MtCEP1) and of Empty Vector (EV) control roots. A Mann-Whitney test was performed between the two genotypes (control empty vector (EV) and 35S:MtCEP1): *, P < 0.05. **B.** Mycorrhizal phenotype of cra2 mutant and Wild-Type (WT) roots. A Student’s t-test was performed between the two genotypes (Wild-Type (WT) and cra2-11 mutant): ***, P < 0.001. Hypha, Intraradical mycelium; IP, Infection Points (fungal entry sites); Arb, Arbuscules; Apex, root tips. Data represent a representative experiment (n = 8 plants per condition) out of three independent experiments. Boxes represent the middle 50% of the values, the median is represented by a horizontal line, and upper and lower quartiles by vertical lines.

### The nodulation-related miR2111 shoot-to-root systemic effector is not regulated by Pi availability nor mycorrhization

Because of the previously characterized conservation of the CEP/CRA2 pathway regulatory function in promoting the two root endosymbioses in legumes, we then wanted to evaluate the potential involvement of the microRNA (miRNA) miR2111 downstream effector that is key to regulate nodulation depending on nitrogen availability^4,15^. This miRNA negatively impacts the transcripts accumulation of the TML1 and TML2 (Too Much Love 1 and 2) genes that act in roots as inhibitors of nodulation initiation^4,16,17^. In M. truncatula roots, the transcripts accumulation of the miR2111, as well as of TML1 and TML2 was not significantly regulated either in response to AMS or to Pi availability (Figure S7). Knowing that in response to nitrogen availability and rhizobium, the miR2111/TML regulatory module is rapidly and strongly regulated through the CRA2 receptor^4^, the Pi availability and AMS expression pattern of the miR2111 and TML genes does not favour a role of this downstream CEP/CRA2-dependent module in the regulation of AMS. This thus suggests that even though there is an evolutionary conservation of the CEP/CRA2 pathway to promote both root endosymbioses, downstream components have likely diverged.

### The CRA2 pathway is required for regulating a subset of Pi-related and AMS-responsive genes, as well as of SL accumulation

To identify downstream molecular targets of the CEP/CRA2 pathway recruited for regulating AMS, a RNAseq analysis was performed to compare the root response to AMS between WT plants and cra2 mutants. A Principal Component Analysis revealed a clearcut discrimination between Non-Mycorrhized (NM) and AMS samples, as well as between cra2 and WT plants (Figure S8). Globally, under low Pi NM conditions promoting AMS, 1218 genes showed an increased expression, and 923 a decreased expression, in cra2 roots compared to WT roots (Table S1). In agreement to the root phenotype of the mutant^3,10^, 22% of the genes more highly expressed in cra2 are associated into lateral root formation (Figure S9A, Table S2^18^). Likewise, the expression of a subset of genes related to auxin transport and responses, cell cycle, and root development, was altered in the cra2 mutant (Figure S9B). In the NM condition, some of these auxin-related genes, and notably MtYUC2 (YUCCA 2), an auxin biosynthesis gene, were previously reported as being similarly induced in cra2 (^19^; Figure S9B). A survey of Gene Ontologies (GO) revealed that GO terms related to detoxification and reactive oxygen species responses were enriched for genes more highly expressed in cra2 mutant roots, while GO terms related to cell wall biogenesis processes were enriched for genes that are more lowly expressed in cra2 (Figure S10), as previously documented^19^. In AMS-colonized roots, 80% and 69% of the genes that were upregulated or downregulated by AMS, respectively, are similarly regulated in cra2 and in the WT (Figure 3A, Figure S11). This correlates with the observation that cra2 mutants engage AMS as in WT plants (Figure 2B), from the infection stage to the propagation of intra-radical mycelium and arbuscule formation, even though its overall colonization rate is reduced. The remaining genes regulated by AMS in the WT and that have lost their regulation in the cra2 mutant show an enrichment for GOs related to amino acids and RNA metabolism, respectively for AMS-upregulated and downregulated genes (Figure S11). Among those, a subset of genes once more related to auxin signalling and transport, cell cycle, and root development are not anymore regulated by AMS in cra2 mutant roots (Figure S9C).

**Figure 3.**
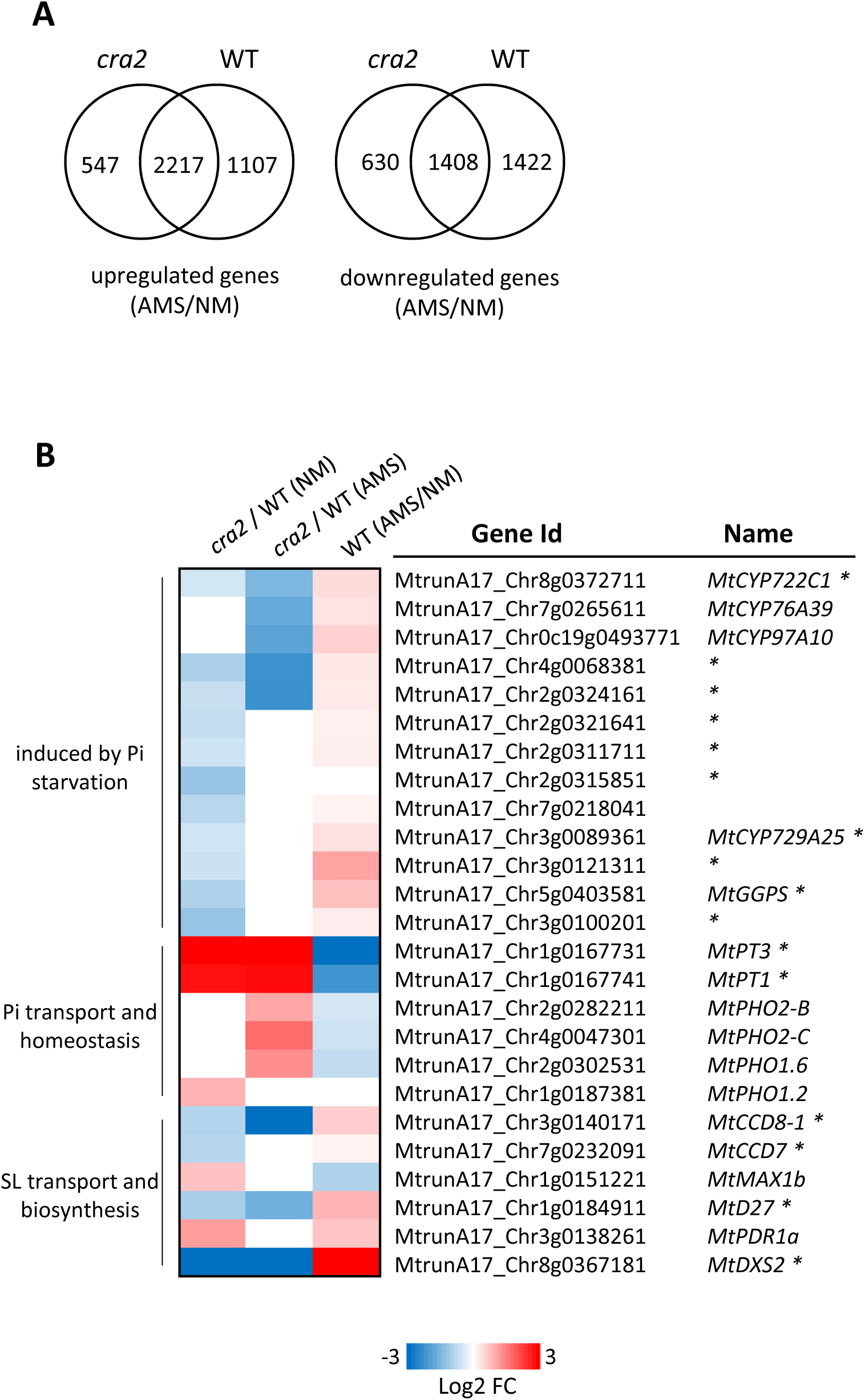
Regulation of selected phosphate- and strigolactone-related genes in Wild-Type versus cra2 mutant roots. **A.** Venn analysis of up- or down-regulated genes (left or right, respectively) in six weeks old Arbuscular Mycorrhizal Symbiotic (AMS) colonized roots relative to Non-Mycorrhized (NM) roots of Wild-Type (WT) or cra2 mutant plants. **B.** Heat map of the expression pattern of strigolactone (SL) biosynthesis and transport, or Pi-starvation response, transport, or biosynthesis genes, misregulated in cra2 mutant roots compared to the WT in NM and/or AMS conditions. Genes followed by an asterisk are also misregulated in 35S:MtCLE53 over-expressing roots^23^ the same way as in cra2 NM roots. Gene annotations from the LeGOO database were used^51^. All displayed fold changes are statistically significant (FDR < 0.05).

Focusing on genes already known to regulate AMS, a subset of these symbiotic genes have a lower expression in cra2 mutant roots compared to the WT already in the NM condition, including MtRAM2 (Reduced Arbuscular Mycorrhization 2), a lipid biosynthetic enzyme required for lipid delivery to the symbiotic fungus^20,21^, and the presumed lipid transporter MtSTR2 (stunted arbuscules 2) proposed to deliver these lipids to the fungus during AMS (^21,22^; Table S3). In addition, the SUNN (Super Numeric Nodules) gene has an increased expression in NM roots in the cra2 mutant compared to the WT (Table S3). SUNN is a Leucine-rich repeat receptor-like protein kinase that was previously shown to repress AMS through the inhibition of the strigolactone (SL) hormonal pathway^23,24^. Moreover, in AMS-colonized roots, SUNN also shows an increased expression in cra2 compared to the WT, while it shows in contrast a repression in WT AMS-colonized roots compared to NM roots (Table S3), suggesting that the CEP/CRA2 pathway may mediate a negative feedback regulation on MtSUNN expression.

A number of genes related to SL transport and biosynthesis were more lowly expressed in roots of cra2 mutant plants grown in the NM condition, compared to WT roots (Figure 3B). Moreover, in the same condition, a subset of genes involved in the Pi-starvation response^25^ were also expressed at lower levels in cra2 mutant roots than in WT roots (Figure 3B). Interestingly, most of these genes show a similarly altered expression pattern in roots overexpressing MtCLE53 (Figure 3B), a CLE peptide acting through SUNN to inhibit SL biosynthesis, and thus AMS^23^. Even more strikingly, this altered cra2 mutant Pi-starvation and SL response was maintained in AMS-colonized roots, such as for the MtCCD8-1 (Carotenoid cleavage dioxygenase 8), MtD27 (DWARF27), and MtDXS2 (1-deoxy-D-xylulose-5-phophate synthase 2) enzymes that act at early (MtDXS2) or late steps (MtCCD8-1, MtD27) of SL biosynthesis (Figure 3B). These results, combined with the previously documented SUNN-mediated inhibition of AMS associated to an altered regulation of SL biosynthesis^23,24^, suggest that SUNN and CRA2 antagonistically regulate AMS by modulating SL biosynthesis. To identify additional downstream targets showing SUNN-dependent and CRA2-dependent antagonistic regulations in the AMS context, we then identified the gene set that is upregulated in cra2 and downregulated in sunn in AMS roots, using the Karlo et al. (2020) dataset^24^. Unexpectedly, genes associated with defence and salicylic acid-mediated gene ontology (GO) terms were significantly enriched (Figure S12A, Table S5). In addition, genes related to Pi transport and metabolism were the most enriched GO terms among the set of genes that are downregulated in cra2 and upregulated in sunn during AMS (Figure S12B, Table S5), further indicating that the antagonistic regulation of AMS by CEP/CRA2 and CLE53/SUNN pathways is tightly linked to the regulation of Pi homeostasis.

As SL production is upregulated in response to Pi starvation conditions^25^, this RNAseq analysis strongly suggests that the cra2 mutant may be impaired in activating a proper Pi-starvation response. In agreement, a subset of genes relative to Pi transport, the Pi transporters 1 (MtPT1) and 3 (MtPT3), showed a higher expression in cra2 in the NM condition, and remained high in AMS roots, while they are in contrast repressed by AMS in the WT (Figure 3B). Interestingly, two of the three paralogs of PHO2 (Phosphate 2), MtPHO2-B and MtPHO2-C^26^, encoding E2 enzymes that are master regulators of Pi acquisition through promoting the ubiquitination and subsequent degradation of Pi transporters, and that are strongly repressed by P-starvation^26,27^, show a higher expression in cra2 mutant AMS roots compared to the WT (Figure 3B). In M. truncatula, the repression of PHO2 expression by AMS was previously proposed to impede the downstream regulation of Pi acquisition in root cortical cells accumulating high Pi amounts due to symbiotic exchanges^28^. Finally, it is also worth mentioning that neither the MtPHR2 (PHOSPHATE RESPONSE 2) or SPX (SYG1/Pho81/XPR1) Pi sensing genes, nor the MtNSP2 (Nodulation Signalling Pathway 2)^29–31^ MtPHR2-downstream target, which were recently shown in M. truncatula as well as in rice^32^ to be both required for the P-starvation activation of SL biosynthesis in response to AMS colonization^25^, are misregulated in the cra2 mutant (Table S1).

As SL biosynthesis genes were lowly expressed in the cra2 mutant, we determined whether SL levels could be affected in cra2 mutant roots grown in Pi-starvation. A SL quantification revealed that didehydro-orobanchol accumulation, also known as medicaol, a potent activator of AMF hyphal branching^33^, was strongly reduced in cra2 mutant roots (Figure 4A). This result corroborates the reduced expression of SL biosynthesis genes in the mutant. As SL are major plant root exudate components activating AMF hyphal branching^34^ and thus allowing AMS^35^, these data suggest that the reduced AMS colonization events observed in cra2 mutant roots may be related to its inability to release sufficient amounts of SL in the rhizosphere. To test this hypothesis, cra2 mutant and WT root exudates were applied to the seeds of the parasitic plant Phelipanche ramosa that require rhizospheric SLs as a germination-promoting signal to efficiently infect their hosts^36^. While WT root exudates, or the GR24 synthetic SL control, triggered the induction of P. ramosa seeds germination, as expected, cra2 mutant root exudates were nearly inactive (Figure 4B and S13), in agreement with previous reports showing that mutants unable to produce SLs are unable to stimulate P. ramosa seed germination^29^. This result indicates that cra2 mutant roots not only accumulate less SLs, but that they are also exuding much less SLs in their rhizosphere. Overall, this could likely explain the low ability of cra2 mutant roots to establish AMS.

**Figure 4.**
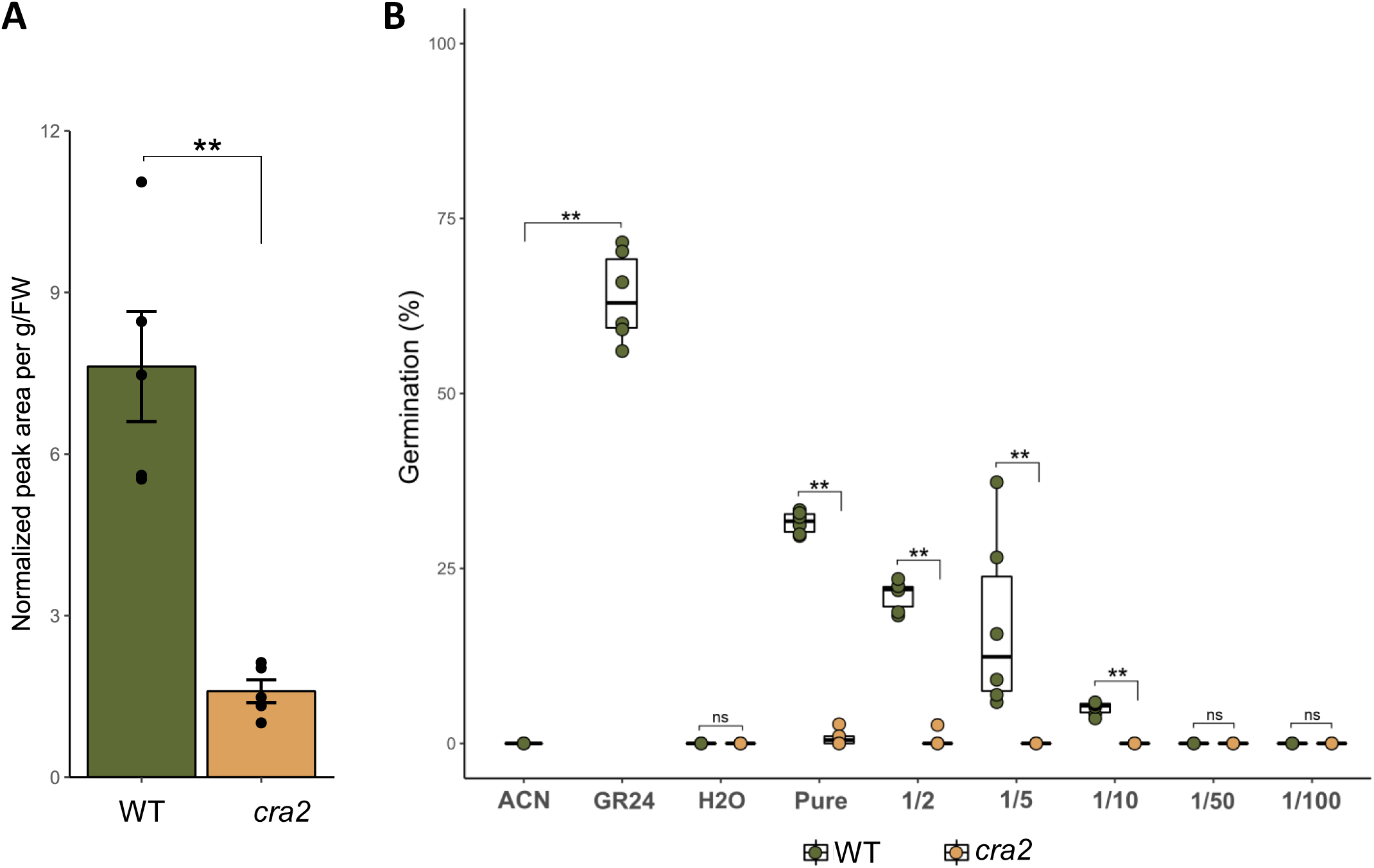
Strigolactones quantification in cra2 mutant roots and P. ramosa seed germination in response to cra2 root exudates. Four-week-old Wild-Type (WT) and cra2 mutant plants were grown under low Pi condition. Five pools of 3-5 root systems were then sampled for didehydro-orobanchol quantification (**A**), or a pool of five plants was transferred for 24h in sterile water to collect the root exudates and perform for P. ramosa seed germination assays (**B**). ACN, acetonitrile 0.001%; GR24, rac-GR24 10^−7^ M; pure, undiluted root exudates; 1/x : serial dilutions of pure root exudates with sterile water. Boxes represent the middle 50% of the values, the median is represented by a horizontal line, and upper and lower quartiles by vertical lines. A Student’s t-test was performed between WT and cra2 plants (A): **, P < 0.01; and a Mann-Whitney between WT and cra2 plants (B): **, P < 0.01.

In the symbiotic nodulation context, the activation of the CEP/CRA2 pathway in low mineral nitrogen conditions that are required for initiating the nitrogen-fixing symbiotic interaction, independently of the rhizobium microsymbiont, revealed its role in determining the root competence to nodulate depending on host plant needs, i.e. only when mineral nitrogen availability is limited^4,17^. Similarly, the activation of the CEP/CRA2 pathway in low Pi conditions independently of AMF colonization likely reflects its role in promoting root competence for initiating AMS depending on host plant metabolic needs. In addition, the transcriptomic and SL quantification data obtained suggest a model where this Pi-starvation promotion of AMS by the CEP/CRA2 pathway would mainly act by regulating the expression of a subset of Pi transport and response genes as well as of SL-biosynthesis genes expression and thus SL accumulation and exudation into the rhizosphere, through a pathway independent of the previously characterized AMS-promoting SPX/PHR2/NSP2 module. SL level reduction in cra2 mutant roots and exudates would thus limit the root competency to engage into AMS.

## Materials & Methods

### Lead Contact and Materials Availability

Further information and requests for resources and reagents should be directed to and will be fulfilled by the Lead Contacts Nicolas Frei dit Frey and Florian Frugier. All unique/stable reagents generated in this study are available from the Lead Contacts without restriction.

### Data Availability

The RNAseq datasets supporting the current study have been deposited in the NCBI/GEO public repository under the accession number XXX (in progress).

### Experimental Model and Subject Details

The Medicago truncatula Jemalong A17^37^ and R108^38^ genotypes, the cra2-1 and cra2-11 mutants that contain respectively a Tnt1 retro-transposon insertion in the leucine-rich repeat region or in the kinase domain encoding region (Key resources table) were used in this study. Seeds were scarified with 95-98% sulfuric acid (Sigma-Aldrich) during eight minutes and washed five times with water. They were then surface sterilized with bleach (9.5%) for one min, washed again six times with sterilized water, and transferred onto 10% w/v water-agar (Sigma # A9799) solid medium plates. The plates were kept in the dark at 4°C for 24 h, and then transferred at 24°C overnight for germination. For contrasting Pi availability experiments, plants were grown in the following different conditions without AMF inoculation: 0.5X Long Ashton medium^39^ modified with low (7.5 μM) or high (2 mM) Pi (NaH_2_P0_4_) in pots containing a mix of zeolite 1.0-2.5 mm and zeolite 0.5-1.0 mm fractions (v/v) as substrate (Symbiom). Plants were grown with a 16h photoperiod and 160 µE light intensity, at 24°C, and with 60% of relative humidity.

For inoculation with AMF, plants were grown in pots containing a mix of zeolite 1.0-2.5 mm and zeolite 0.5-1.0 mm fractions (v/v) as substrate (Symbiom), watered twice a week with 0.5X Long Ashton medium^39^ containing 7.5 μM NaH PO , and inoculated directly after germination with 300 spores of the Rhizophagus irregularis DAOM 197198 strain (Agronutrition; Key resources table). Roots were harvested six weeks after AMF inoculation, washed three time with water, before staining (see below).

For root exudate collection, plants were grown for four weeks under the low Pi condition described above. Five plants per genotype were removed from the substrate, their root system rinsed with water, and plants were then transferred in 50mL Falcon tubes with 20mL of water and shoots outside the open tube. After 24h, plants were removed and the water containing root exudates was sterilized with a 0.2 µm filter and further used, pure or diluted with water, for germination assays with Phelipanche ramosa^40^ seeds (Key resources table).

## Method Details

### AMF colonization phenotyping

Whole plant root systems were collected six weeks after AMF inoculation, washed with water, treated with 10% KOH for two days at room temperature, washed again with water, and finally stained with an ink solution (5% Schaeffer black ink, 95% acetic acid; Key resources table) for 5min at 95°C^41^ . Stained roots were then washed again with water, and cleared with 50% ethanol.

The quantification of mycorrhizal colonization was performed using the grid-intersect method^42^. Entire root systems were cut into small fragments (∼2 cm) and randomized under the microscope. A minimal WT plant root colonization of approximately 50% was validated in all independent experiments before proceeding to the detailed quantifications. At least 400 root intersections were observed for each plant root system (n = 8-10 per genotype) to quantify root apices, fungal infection points at the root epidermis (hyphopodia), as well as arbuscules and intraradical mycelium colonization of the root cortex. Observations were performed with a S6E microscope (Leica microsystems).

#### Agrobacterium rhizogenes-mediated transformation

The p35S:CEP1 construct, as well as the control empty vector pK7WG2D^43^, were generated in Imin et al. (2013)^1^; Key resources table), and used for root transformation using the Agrobacterium rhizogenes Arqua1 strain (Key resources table), as described in Boisson Dernier et al. (2001)^44^. The p35S:CEP2 construct was generated in the p2L-50507 plasmid^45^, and the p2L-50507 plasmid was used as control empty vector. Composite plants, with WT shoots and transgenic roots, were grown in vitro on Fahraeus medium^46^ plates and inoculated with AMF as described above.

### RNA extraction and cDNA synthesis

Total RNAs were extracted using the Quick-RNA Miniprep kit (Zymo, Key resources table), from roots which were collected on plants grown in low versus high Pi conditions four weeks after germination, or six weeks after inoculation with the AMF. In all cases, pools of three plants were used as biological replicates. The DNAse treatment was performed with a RNAse-free DNAse1 following the manufacturer instructions. cDNAs were then obtained with the Reverse Transcriptase Superscript III

(Invitrogen; 200U/μL; Key resources table) following manufacturer instructions. A Stem-Loop Reverse Transcription (RT) was performed to amplify selected mature 21 base pairs miRNAs thanks to dedicated adapters (Table S4), as described in Proust et al. (2018)^47^.

### Real-Time PCR analysis

Gene expression was analysed by quantitative RT-PCR (qRT-PCR) on a LightCycler96 apparatus (Roche) using the Light Cycler 480 Green I Master mix (Roche, Key resources table) and specific primers to amplify the genes of interest (listed in Table S4). Forty-five cycles of amplification (15 sec at 95°C, 15 sec at 60°C, 15 sec at 72°C) were performed, with a final melting curve step (from 60 to 95°C) to assess primers specificity. All primers used were selected with an efficiency of at least 90%. Mycorrhizal and Pi starvation marker genes were respectively described in Feddermann et al. (2008)^48^ and Branscheid et al. (2010)^28^. Gene expression normalizations were performed with the reference genes MtActin11 (MtACT11) and MtRBP1, and with the miR162 for mature miRNAs, based on previous studies^4,49^. As the normalization with MtACT11 or MtRBP1 gave similar results, only the normalization with MtACT11 is shown in figures.

### RNAseq transcriptomic analysis

To perform differential analyses, htseq-counts files (Genewiz), mapped onto the v5 version of the M. truncatula genome^37^, were analysed with the R software using the EdgeR package version 3.24.3^50^(Key resources table). Gene annotations from the LeGOO database were used^51^ (Key resources table). Genes which did not have at least one read after a count per million in at least one half of the samples were discarded at first. Then, raw counts were normalized using the Trimmed Mean of M-values method. The count distribution was modelled with a negative binomial generalized linear model where the condition, the genotype and the interaction between conditions and genotypes were taken into account, and where the dispersion was estimated with the EdgeR method^50^. A likelihood ratio test was performed to evaluate the effect of the condition in each genotype, or the effect of the genotype in each condition. Raw P values were adjusted with the Benjamini–Hochberg procedure to control the False Discovery Rate (FDR). A gene was declared as differentially expressed if its adjusted P value was ≤ 0.05.

GO term enrichments were determined using the ShinyGO web tool^52^ (Key resources table) with default parameters, and Venny^53^ (Key resources table) was used to perform Venn diagrams.

### Extraction of Strigolactones from roots

For the SL content analysis, 500 mg of ground roots (six-week-old plants grown in low Pi) were transferred to 10 mL glass vials. SLs were extracted with 2 mL of ethyl acetate containing 10^−8^ M GR24 (Chiralix, Key resources table) as an internal standard (final concentration 10^−7^ M). Vials were vortexed, sonicated for 20 s (Branson Ultrasonics sonication bath; Emerson Automation Solutions, Busseno, Italy), and extracted at 4°C overnight on an orbital shaker. Samples were centrifuged for 10 min at 2500 g at room temperature. Organic phases were transferred to 4 mL glass vials, and the solvent was evaporated using a speed vacuum system (SPD121P; ThermoSavant, Hastings, UK). SL from roots were quantified according to Liu et al. (2011)^54^.

### Detection and Quantification of Strigolactones

Sample residues were dissolved in 100 µL of acetone/water (30:70, v/v) and filtered through a 0.45 mm Minisart SRP4 filter (Sartorius, Goettingen, Germany). The SL analysis was performed by comparing retention times and mass transitions as previously described by Volpe et al. (2023)^55^, with modifications. A Waters Xevo TQs mass spectrometer with an electrospray ionization source coupled to an Acquity UPLC system (Waters, Milford, USA) was used. Chromatographic separations were conducted on an Acquity UPLC BEH C18 column (100 mm, 2.1 mm, 1.7 mm; Waters, USA) using an acetonitrile/water (0.1% formic acid) gradient. The column was operated at 50°C with a flow rate of 0.5 mL/min. The gradient started at 5% (v/v) acetonitrile for 0.67 min, increased to 27% at 5 min, and to 65% at 8 min. The gradient then increased to 90% acetonitrile in 0.5 min, was maintained for 1 min, before returning to 5% for 2.5 min. The sample injection volume was 5 µL. The mass spectrometer was operated in the positive electrospray ionization mode, with cone and desolvation gas flows set to 150 and 1000 l/h, respectively. The capillary voltage was 2.5 kV, source temperature 150°C, and desolvation temperature 500°C. Argon was used for fragmentation by collision-induced dissociation. Multiple Reaction Monitoring (MRM) was used for quantification. MRM transitions, cone voltage, and collision energy for didehydroorobanchol and GR24 were described by Volpe et al. (2023)^55^.

#### P. ramosa germination assays

P. ramosa seeds (Key resources table) were surface-sterilized in a 2.4% sodium hypochlorite solution for 4 min under gentle stirring, then rinsed three times for 5 min with sterile distilled water using a cell strainer (EASYstrainer 40 µm, Greiner). Seeds were suspended at a final concentration of 5 mg/mL in a conditioning solution (1 mM 4-(2-hydroxyethyl)-1-piperazineethanesulfonic acid (HEPES), pH 7.5 adjusted with KOH; PPM 0.1% v/v) and incubated in the dark at 21°C for 14 days, as described in Lechat et al. (2015)^56^. Germination was induced in 96-well plates by adding 50 µL of serial dilutions of fresh M. truncatula root exudates to 50 µL of conditioned seeds (∼75 seeds per well). Water, and a 10^−7^ M rac-GR24 treatment (in 0.001% acetonitrile; Chiralix, Key resources table) served as negative and positive controls, respectively. After six days of incubation at 21°C in the dark, seeds were stained with 10 µL of 5 g/L methylthiazolyldiphenyl-tetrazolium bromide (MTT; Sigma-Aldrich) to facilitate counting^57^. The germination rate was assessed one day later under a binocular magnifier (Leica MZ10F).

### Wheat Germ Agglutinin (WGA) Alexafluor-488 staining and confocal microscopy

For WGA AlexaFluor-488 staining, six-week-old mycorrhized M. truncatula A17 and cra2-11 root systems were incubated in a 10% KOH solution for 48 h at room temperature. Following this treatment, M. truncatula roots were then incubated with 0.1% Triton X-100 for 1 h, and finally in 1 μg/mL WGA Alexafluor-488 (Key Resource Table) solution diluted in PBS 1x at 4°C for 48 hours. The roots were kept in the dark, wrapped in a paper foil, and stored at 4°C before imaging. Stained roots were observed with an upright laser scanning confocal microscope (SP8 Leica Microsystems). WGA-Alexafluor-488 was detected using a 488 nm laser and the emitted light was collected under a 498-550 nm window.

### Statistical Analyses

Statistical analyses were performed with the XLSTAT software or using dedicated R scripts (Key resources table). The normal distribution of experimental values was determined with a Shapiro test (P < 0.05). Statistical difference between groups (low Pi versus high Pi, WT versus cra2 mutant, empty vector versus 35S:MtCEP1/MtCEP2 roots) was then determined using a Student’s t test when data were normally distributed, or a Mann-Whitney test when data were not normally distributed.

### Phylogenetic analyses

The phylogenetic analysis of CEP and PIP pre-proprotein sequences, recovered from the de Bang et al. (2017) study^58^, were aligned using the MAFFT^59^ (Multiple sequence Alignment based on the Fast Fourier Transform; Key resources table) method using default parameters. The tree was then constructed by average linkage using the unweighted pair group method with arithmetic means, and displayed using the MAFFT phylo.io tool^60^.

## Acknowledgements

Work in the FF laboratory was supported by the “Centre National de la Recherche Scientifique” (CNRS), the “SYMPA-PEP” project from the French “Agence Nationale de la Recherche” (ANR), and the “Ecole Universitaire de Recherche” (EUR) “Saclay Plant Science” (SPS). Work in the NFdF laboratory was supported by the CNRS, the “SYMPA-PEP” ANR project, the “Fédération de Recherche Agrobiosciences, Interactions et Biodiversité” (FRAIB), and the University Paul Sabatier (UPS) of Toulouse. We thank Jean-Bernard Pouvreau (Nantes University, US2B) for the gift of P. ramosa seeds.

## Authors Contributions

FF and NFdF designed the study; LP, JTS, TR, WK and NFdF performed experiments and analysed the data; FF analysed the data; FF and NFdF wrote the manuscript, with contributions from JTS and LP.

---

**Figure S1.**
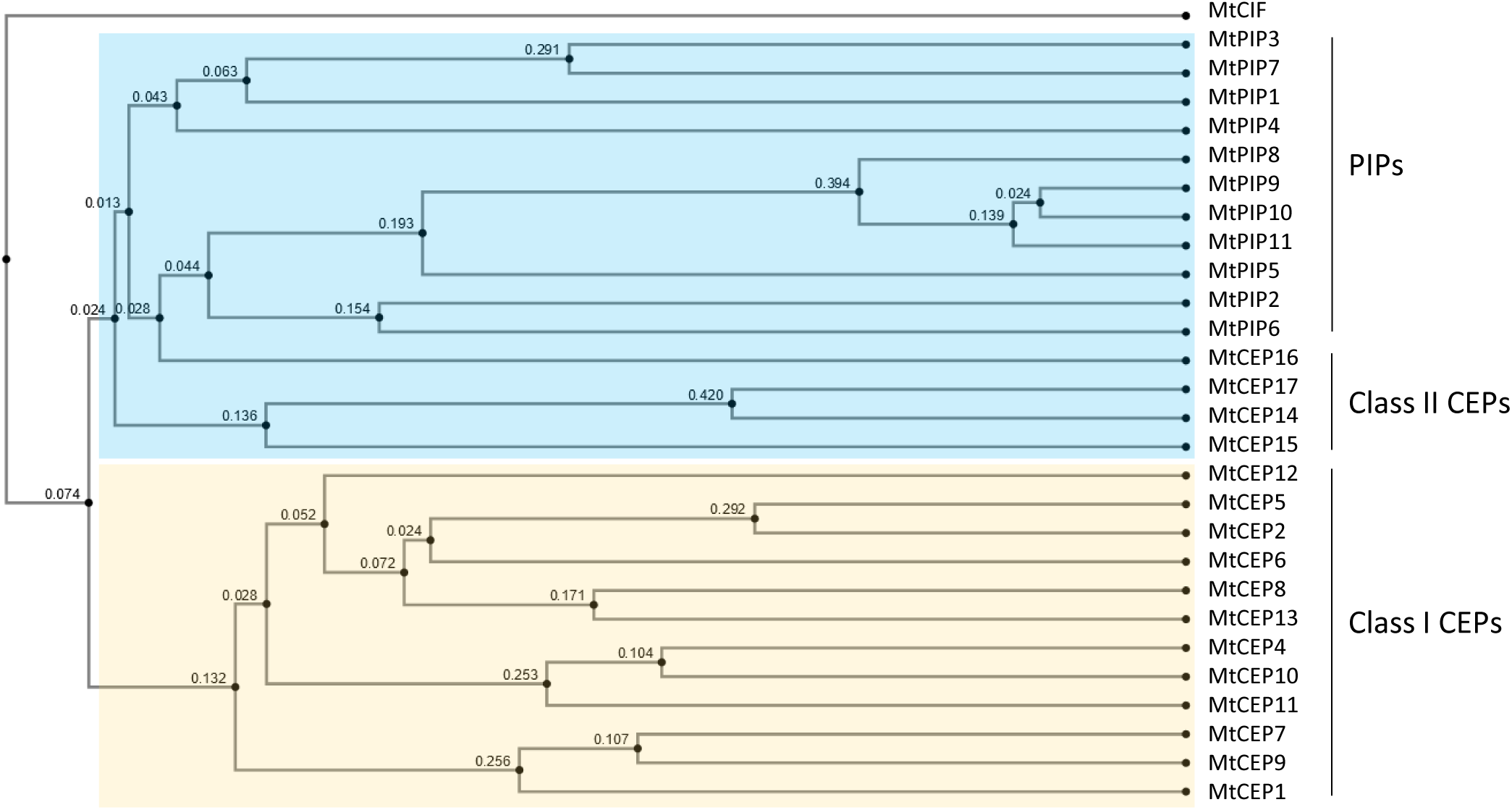
Phylogenetic analysis of Medicago truncatula PIP and CEP signalling peptide encoding genes. The coding sequences of PIP and CEP pre-proproteins^58^ were aligned using the MAFFT software^59^ using default parameters, and the tree was constructed by average linkage using the unweighted pair group method with arithmetic means, and displayed using the MAFFT phylo.io tool^60^. MtCEP10a and MtCEP10b have identical sequences and were then grouped together under the name MtCEP10. MtCEP19 was excluded from the analysis because it is absent from the v5 Medicago genome version^37^, and MtPIP13 was removed because its sequence is truncated at the N-terminus. Class II CEPs and PIPs form a monophyletic group, highlighted in blue, while Class I CEPs, highlighted in yellow, are phylogenetically distant and form and independent group.

**Figure S2.**
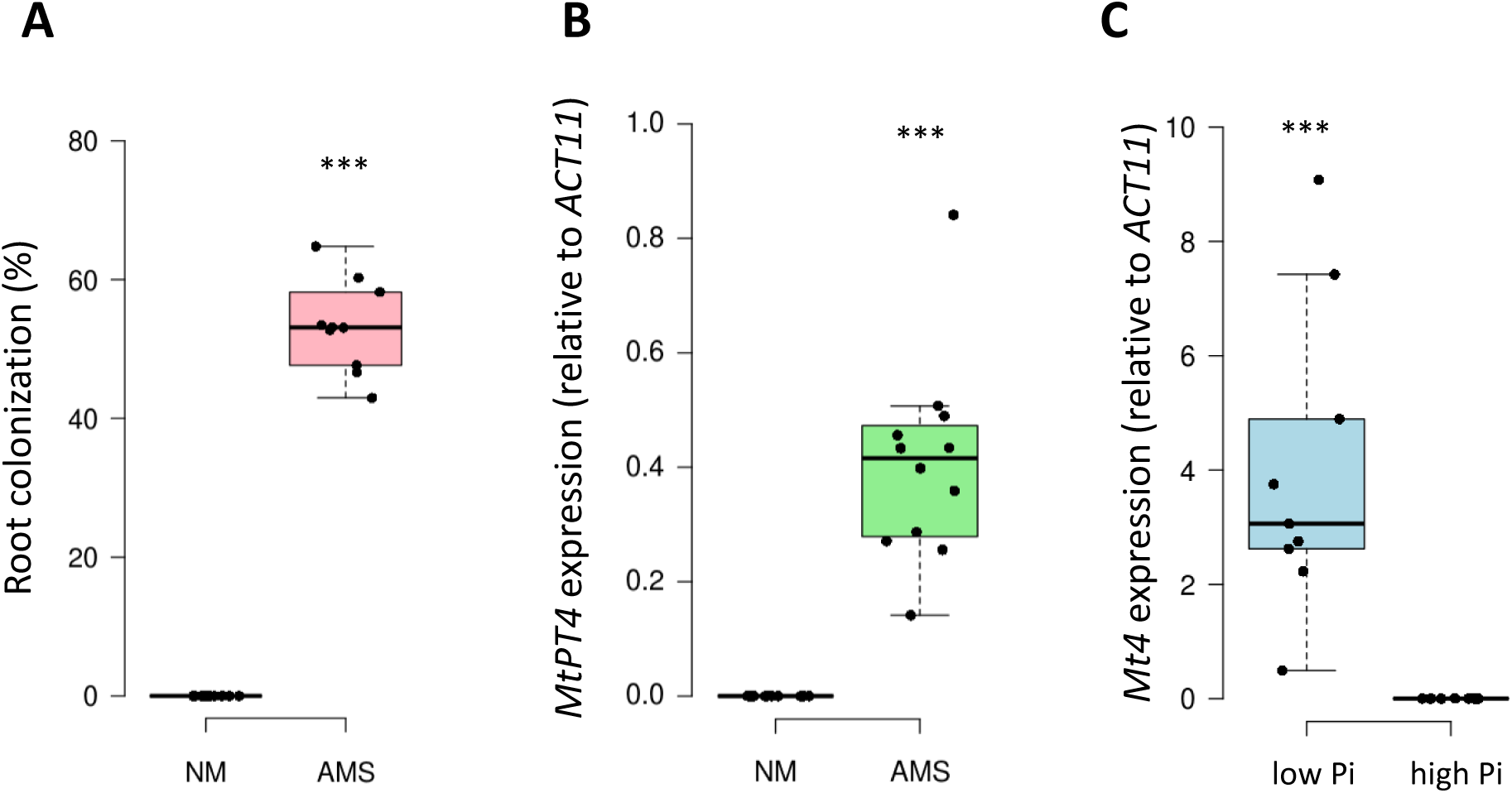
Mycorrhizal colonization, and expression of mycorrhizal and Pi availability markers, in plants used to study MtCEP gene expression (related to Figure 1) **A.** Wild-Type (WT) plants were grown with Arbuscular Mycorrhizal Symbiotic fungi (AMS), or without (NM, Non-Mycorrhized). Plants were used to perform the MtCEP gene expression analyses shown in the Figure 1A. **B.** Regulation of the AMS marker MtPT4. WT plants were grown with Arbuscular Mycorrhizal Symbiotic fungi (AMS) or without (NM, Non-Mycorrhized). **C.** Regulation of the Pi availability marker Mt4. WT plants were grown under low Pi or high Pi conditions. In **B-C**, gene expression values were normalized to the MtACT11 gene. In **A-C**, Data from two independent biological experiments are shown (n=5 pools of 3 plants per experiment). Boxes represent the middle 50% of the values, the median is represented by a horizontal line, and upper and lower quartiles by vertical lines. A Mann-Whitney test was performed between the two conditions (AMS and NM (A, B), or low Pi and high Pi (**C**)) : ***, P < 0.001.

**Figure S3.**
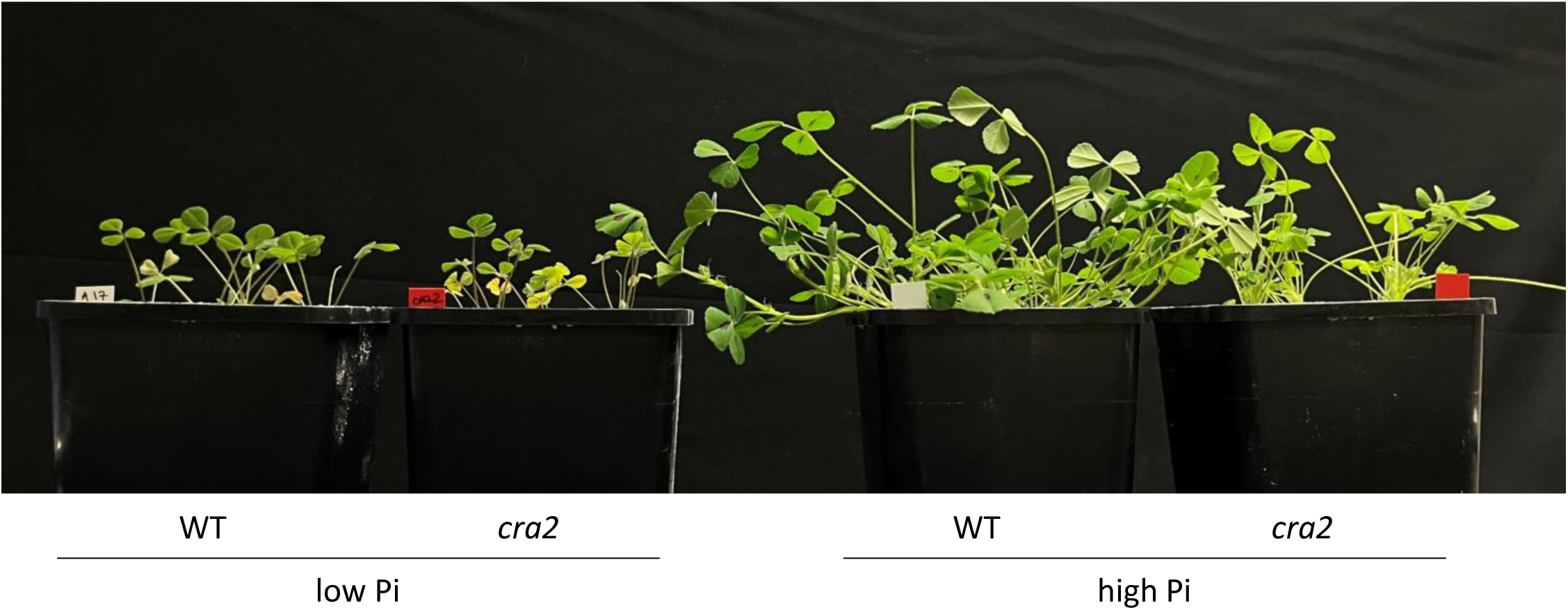
Representative images of Wild-Type and cra2 mutant plants grown under a low Pi or high Pi condition (related to Figure 1B) Representative pictures of Wild-Type (WT) and cra2 mutant plants after 4 weeks of growth under a low (7.5 µM) or high Pi (2 mM) concentration.

**Figure S4.**
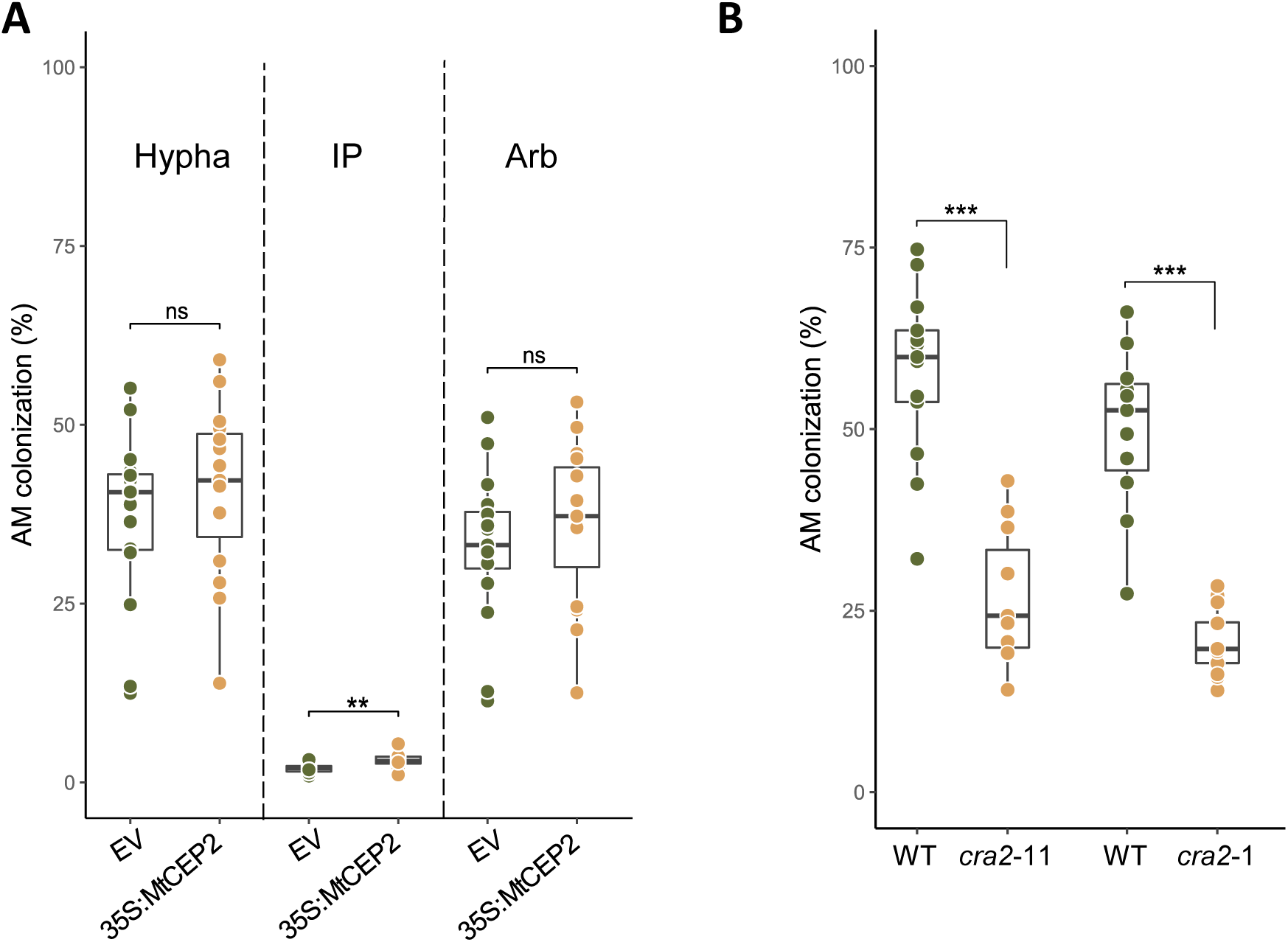
Mycorrhizal phenotype of MtCEP2 overexpressing plants (related to Figure 2) and of cra2-1 and cra2-11 mutant alleles, respectively in the R108 and A17 genotypes. **A.** Mycorrhizal phenotype of MtCEP2 overexpressing (35S:MtCEP2) or Empty Vector (EV) roots. A Mann-Whitney test was performed between the two genotypes (control empty vector (EV) and 35S:MtCEP2): **, P < 0.01. **B.** Mycorrhizal phenotype of cra2-1 mutant or Wild-Type (WT, R108) roots, and cra2-11 mutant or Wild-Type (WT, A17) roots. A Student’s t-test was performed between the Wild-Type (WT) and the cra2-1 mutant genotypes, and a Mann-Whitney test between the WT and the cra2-11 mutant genotypes: ***, P < 0.001. Data represent a representative experiment (n = 8 plants per condition) out of three independent experiments. Boxes represent the middle 50% of the values, the median is represented by a horizontal line, and upper and lower quartiles by vertical lines.

**Figure S5.**
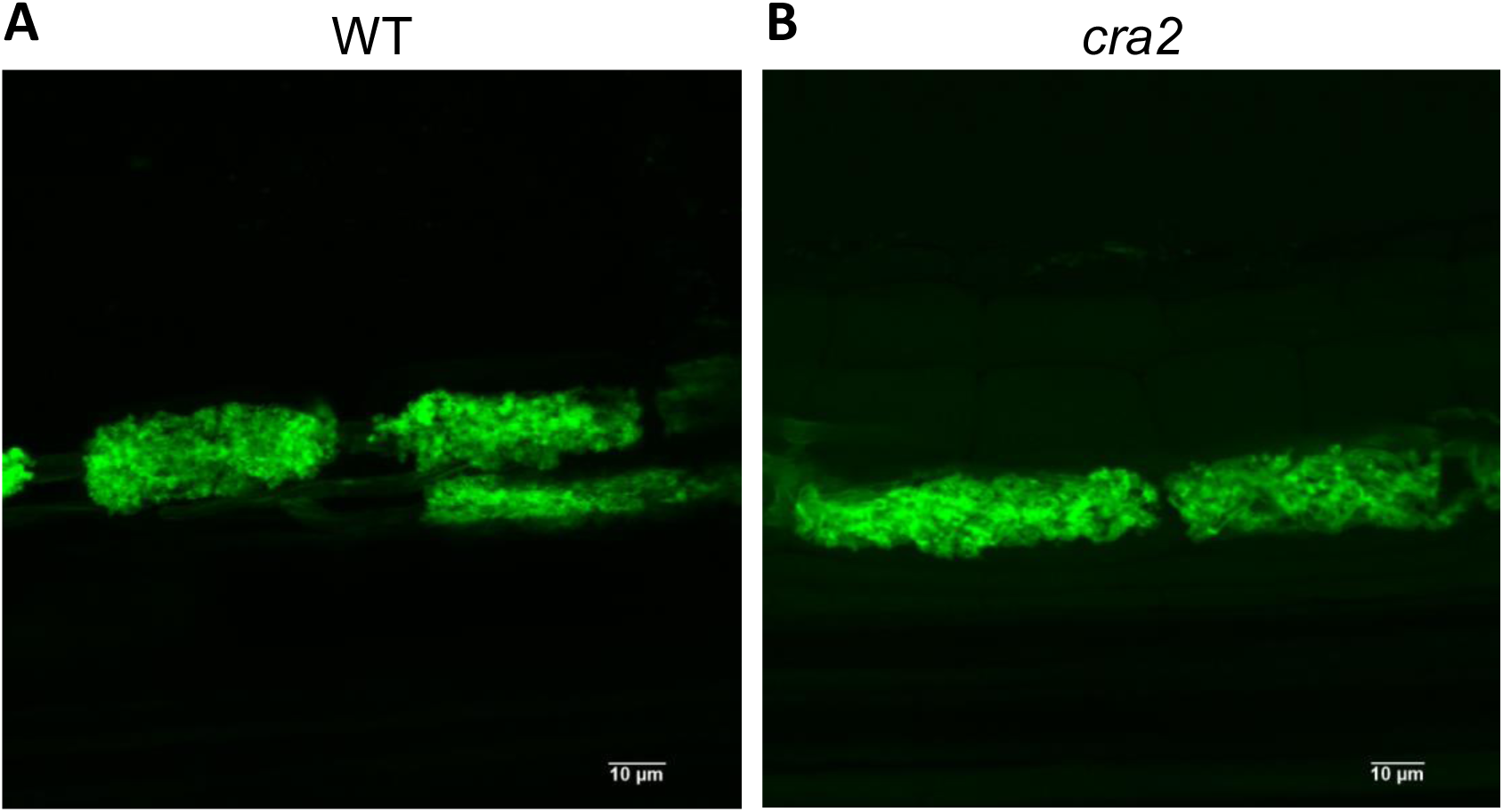
Representative images of WGA-Alexafluor488 stained arbuscules in Wild-Type and cra2 mutant plants (related to Figure 2B) Six-week-old mycorrhized plants were stained with WGA-AlexaFluor488 to highlight fungal arbuscular structures. Ten arbuscules from five plants generated in two independent experiments were observed, and representative images are shown. No major arbuscule developmental phenotype was detected when comparing the two genotypes. Scale bar = 10 µm.

**Figure S6.**
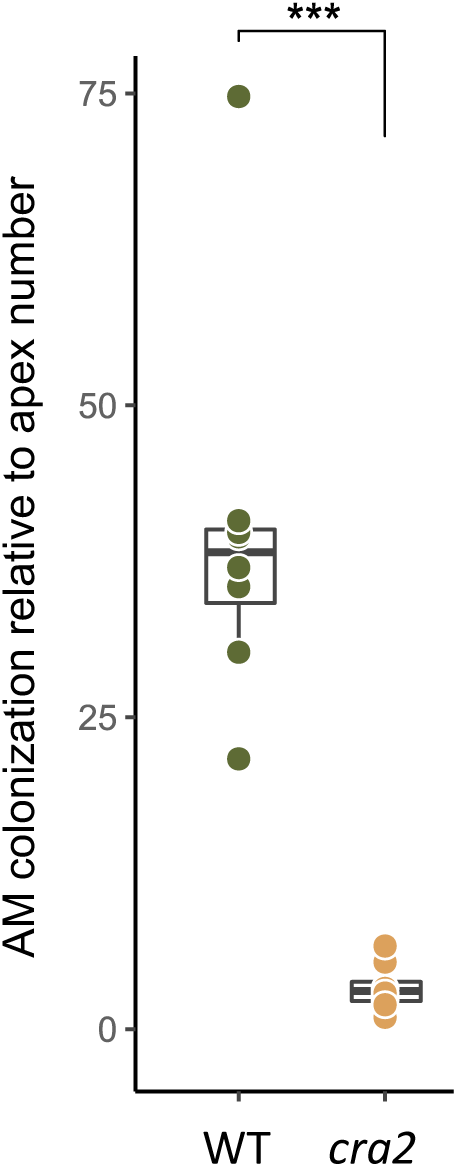
Mycorrhizal colonization of WT and cra2 mutant plants, relative to the percentage of lateral root apices (related to Figure 2B) Mycorrhizal phenotype of cra2 mutant and Wild-Type (WT) roots. The percentage of hyphal colonisation was normalised to the percentage of apices in each root system. A Mann-Whitney test was performed between the two genotypes (Wild-Type (WT) and cra2 mutant): ***, P < 0.001. Hypha, Intraradical mycelium; IP, Infection Points (fungal entry sites); Arb, Arbuscules. Data represent a representative experiment (n = 8 plants per condition) out of three independent experiments. Boxes represent the middle 50% of the values, the median is represented by a horizontal line, and upper and lower quartiles by vertical lines.

**Figure S7.**
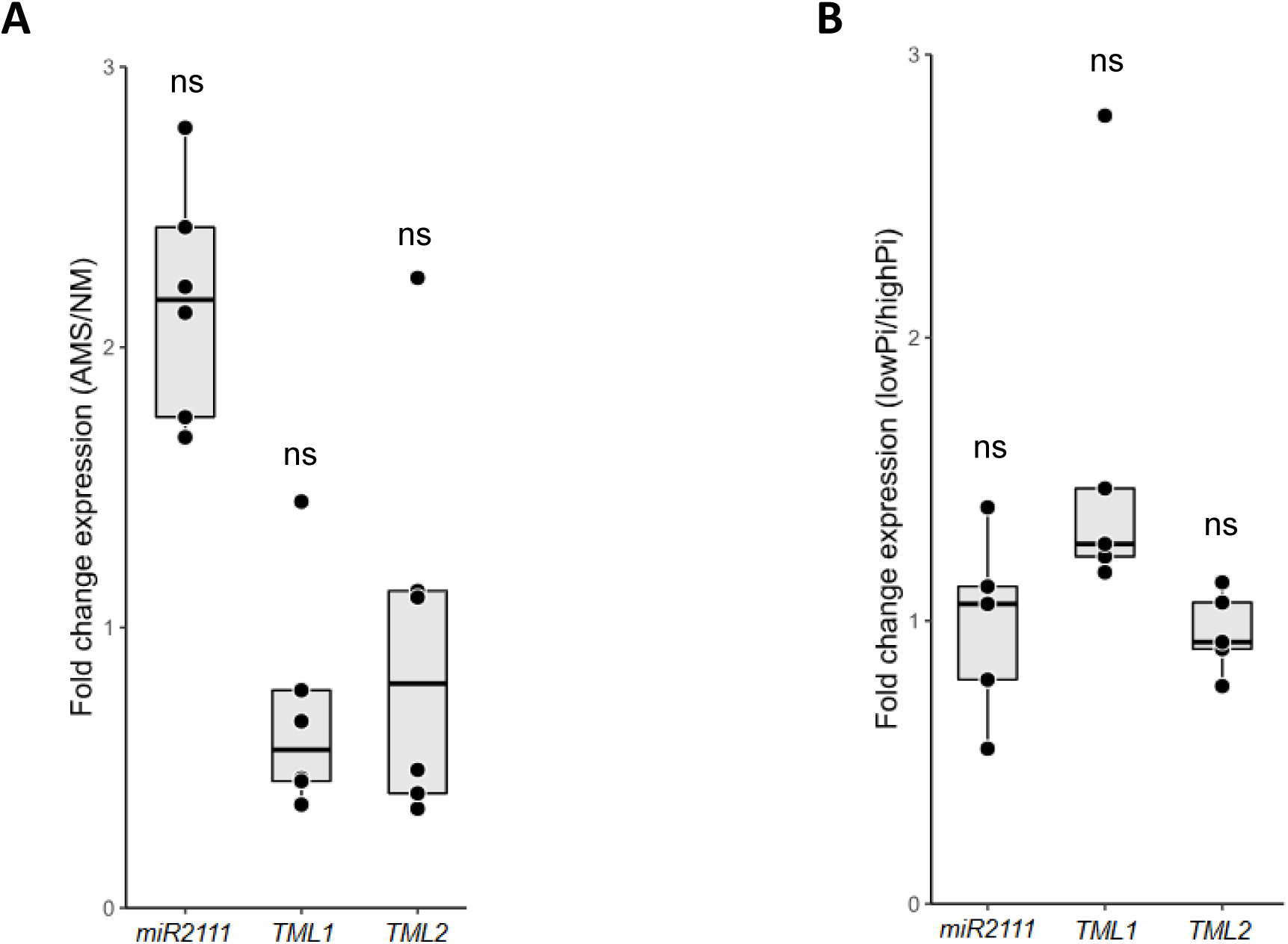
Regulation of the miR2111 / TML module in response to mycorrhization and Pi availability. **A.** miR2111, MtTML1 and MtTML2 transcripts accumulation with Arbuscular Mycorrhizal Symbiotic (AMS) fungi or without (NM, Non-Mycorrhized). **B.** miR2111, MtTML1 and MtTML2 transcripts accumulation in low versus high Pi conditions. Data represent a representative experiment (n = 5 plants per condition) out of three independent experiments. Boxes represent the middle 50% of the values, the median is represented by a horizontal line, and upper and lower quartiles by vertical lines. A Mann-Whitney test was performed between the two conditions (AMS and NM in (**A**), or low Pi and high Pi in (**B**)). ns, no significant difference at P < 0.05 was detected.

**Figure S8.**
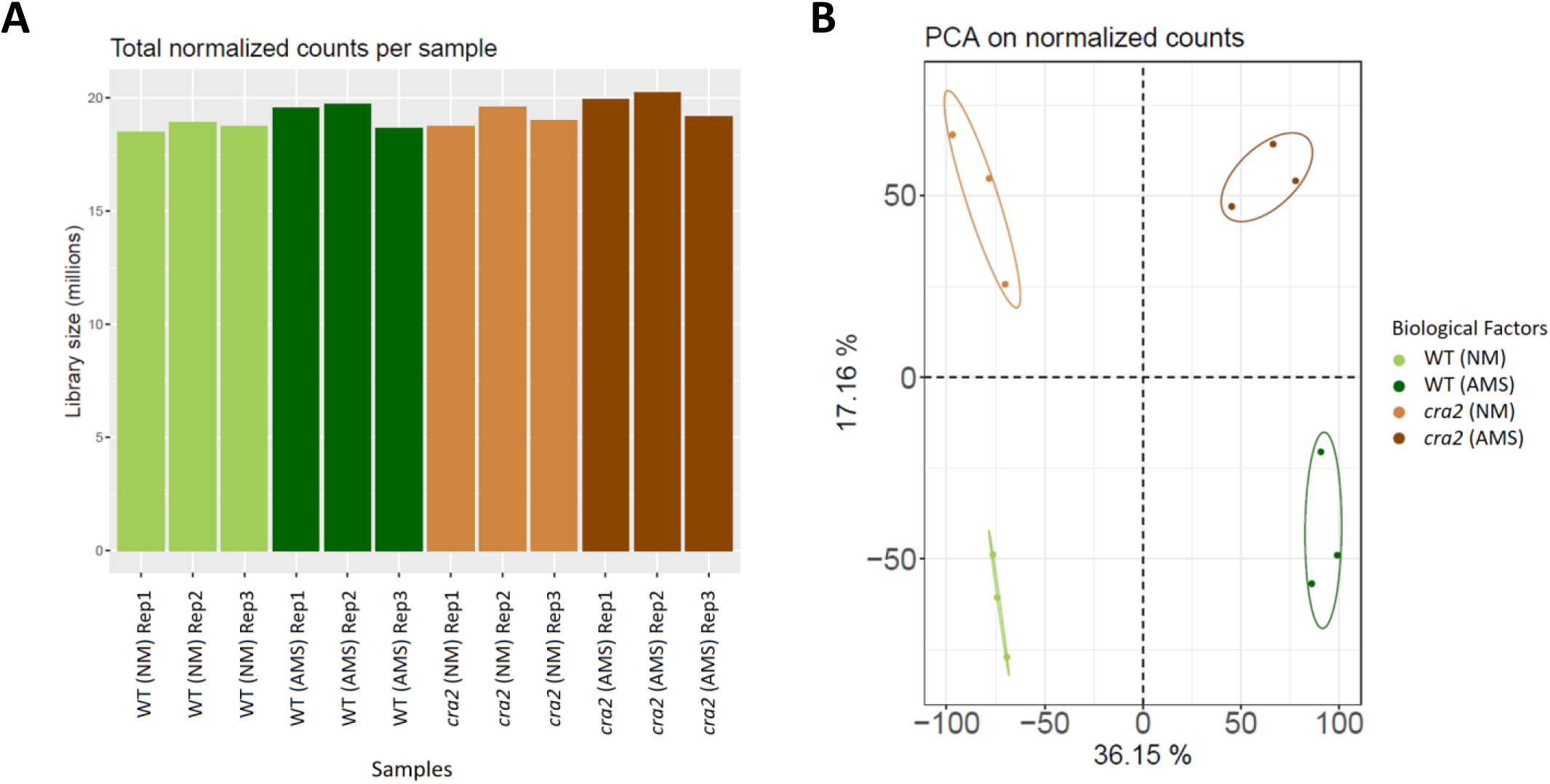
Quality control of the RNAseq analysis (related to Figure 3) **A.** Total normalized counts per sample (n=3 pools of 3 plants) of RNAseq datasets from cra2 and Wild-Type (WT) roots of plants grown with Arbuscular Mycorrhizal Symbiotic fungi (AMS) or without (NM). **B.** Principal component analysis (PCA) on normalized counts, allowing separating the two genotypes and the two treatments.

**Figure S9.**
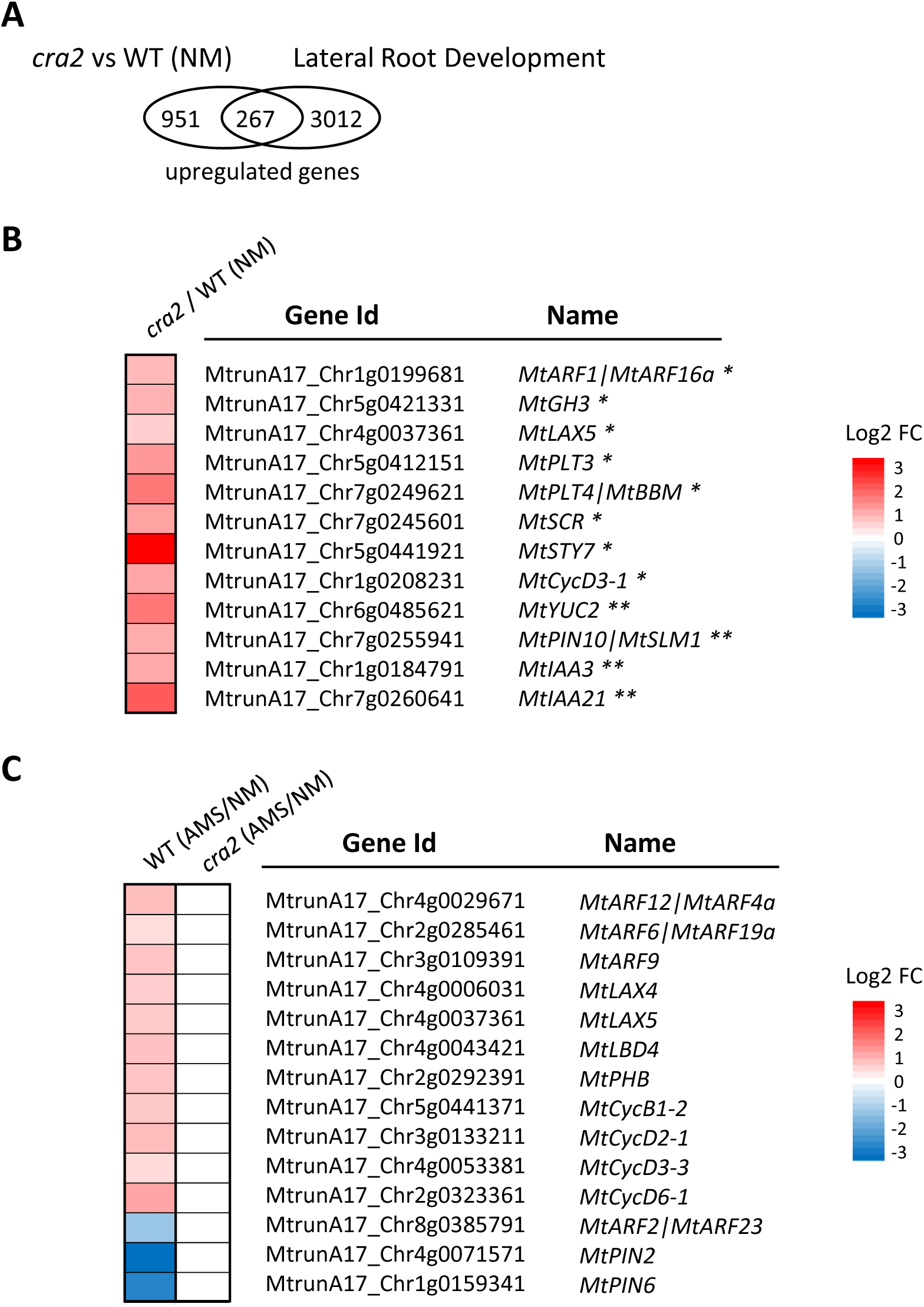
Selected auxin-related, cell cycle-related, and lateral root development-related genes deregulated in cra2 mutant roots (related to Figure 3) **A.** Venn diagram of differentially expressed genes from the RNAseq dataset that are both upregulated in cra2 mutant roots in Non-Mycorrhized (NM) conditions and during lateral root development, based on the Schiessl et al. (2019)^18^ RNAseq. Genes were considered as upregulated during lateral root development if upregulated in at least one time point of the experiment performed by Schiessl et al. (2019)^18^. **B.** Heatmap of a selection of genes of the Venn diagram intersection shown in A and related to auxin signalling and transport, cell cycle, and root development, indicated with *. Auxin-related genes previously identified in cra2 mutant roots generated by CRIPSR-Cas9 in the R108 M. truncatula genotype^19^ are indicated with **. All displayed fold changes are statistically significant (FDR < 0.05). **C.** Heatmap of genes related to auxin signalling and transport, cell cycle, and root development that are up- or down-regulated in Wild-Type (WT) roots in Arbuscular Mycorrhizal Symbiotic (AMS) conditions but unaltered in cra2 mutant roots in the same conditions. All displayed fold changes are statistically significant (FDR < 0.05). Gene annotations from the LeGOO database were used^51^.

**Figure S10.**
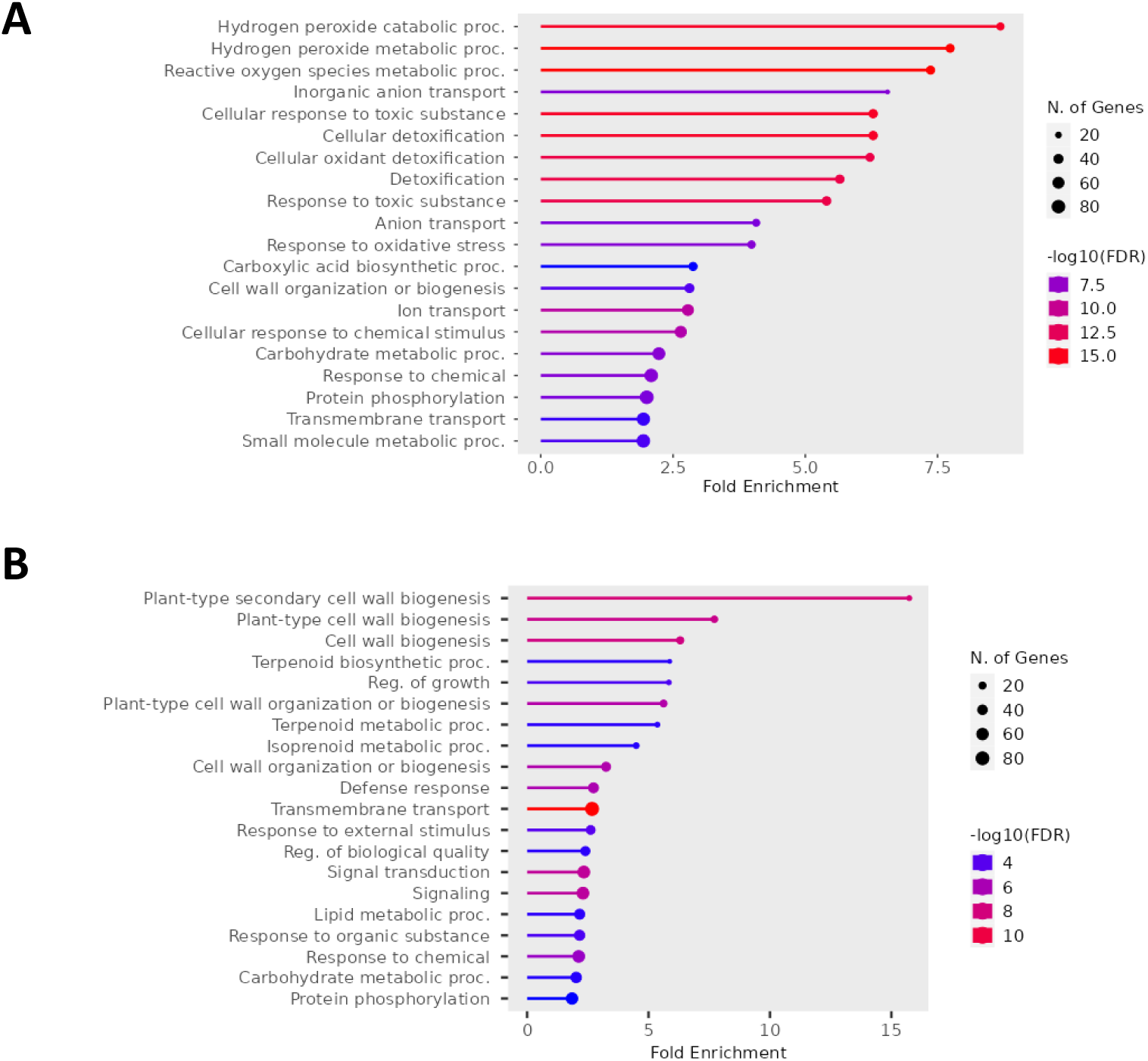
Gene Ontology (GO) enrichments of genes differentially expressed in cra2 mutant roots in Non-Mycorrhizal conditions (related to Figure 3) **A.** Top 20 of enriched “Biological Process” GO terms (hypergeometric distribution test, FDR < 0.05) observed among the 1218 genes upregulated in Non-Mycorrhized (NM) cra2 mutant roots versus NM Wild-Type (WT) roots, from the RNAseq dataset. **B.** Top 20 of enriched “Biological Process” GO terms (hypergeometric distribution test, FDR < 0.05) observed among the 923 genes downregulated in NM cra2 mutant roots versus NM WT roots, from the RNAseq dataset.

**Figure S11.**
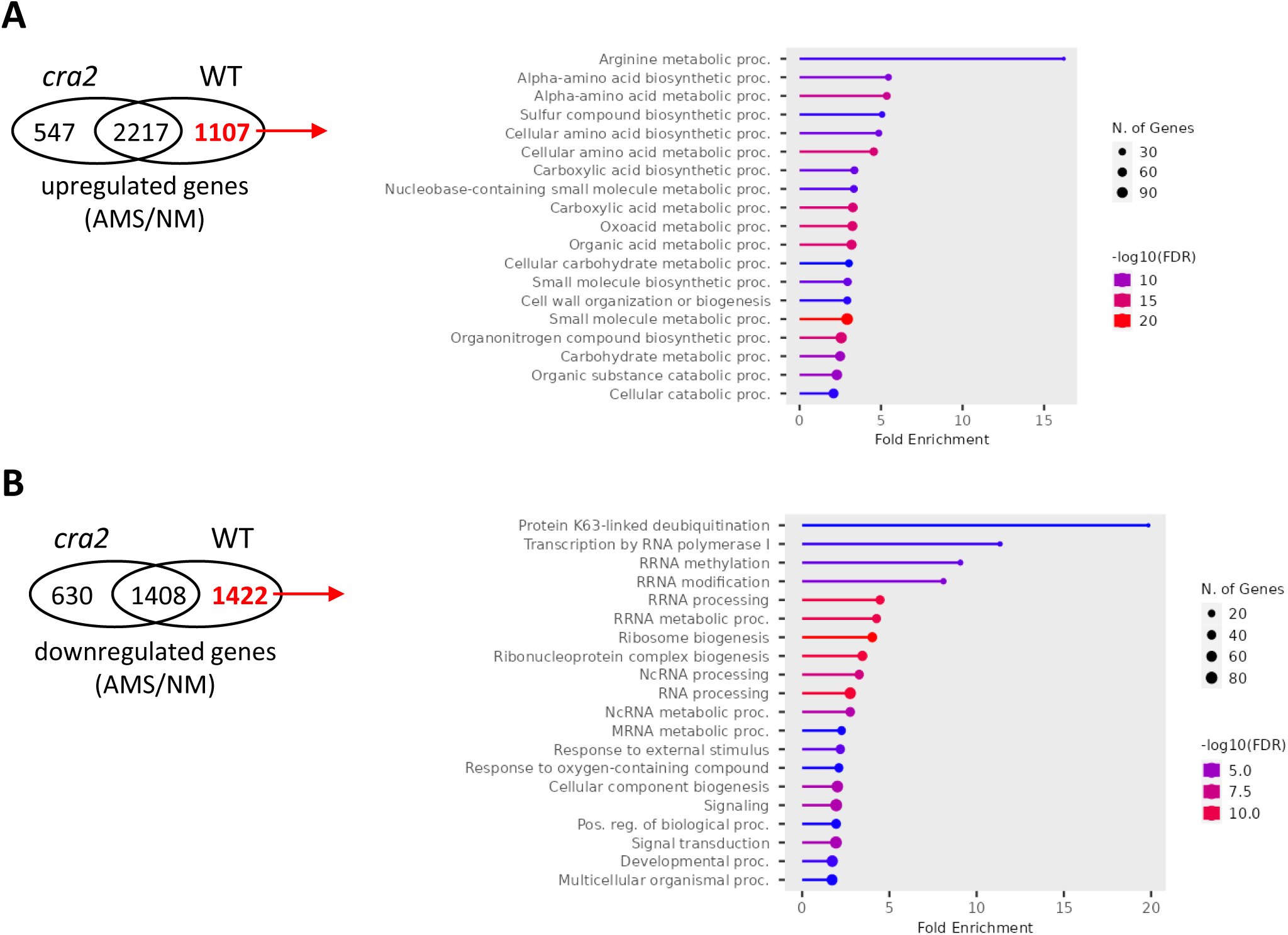
Gene Ontology (GO) enrichments of genes differentially expressed in Wild-Type mycorrhized roots but not in cra2 mutant roots (related to Figure 3) **A.** Top 20 of enriched “Biological Process” GO terms (hypergeometric distribution test, FDR < 0.05) observed among the 1107 genes upregulated in the Wild-Type (WT) in Arbuscular Mycorrhizal Symbiotic fungi (AMS) inoculated versus Non-Mycorrhized (NM) roots, but not upregulated in cra2 mutant roots in the same conditions, from the RNAseq dataset. **B.** Top 20 of enriched “Biological Process” GO terms (hypergeometric distribution test, FDR < 0.05) observed among the 1422 genes downregulated in the WT in AMS versus NM roots, but not downregulated in cra2 mutant roots in the same conditions, from the RNAseq dataset.

**Figure S12.**
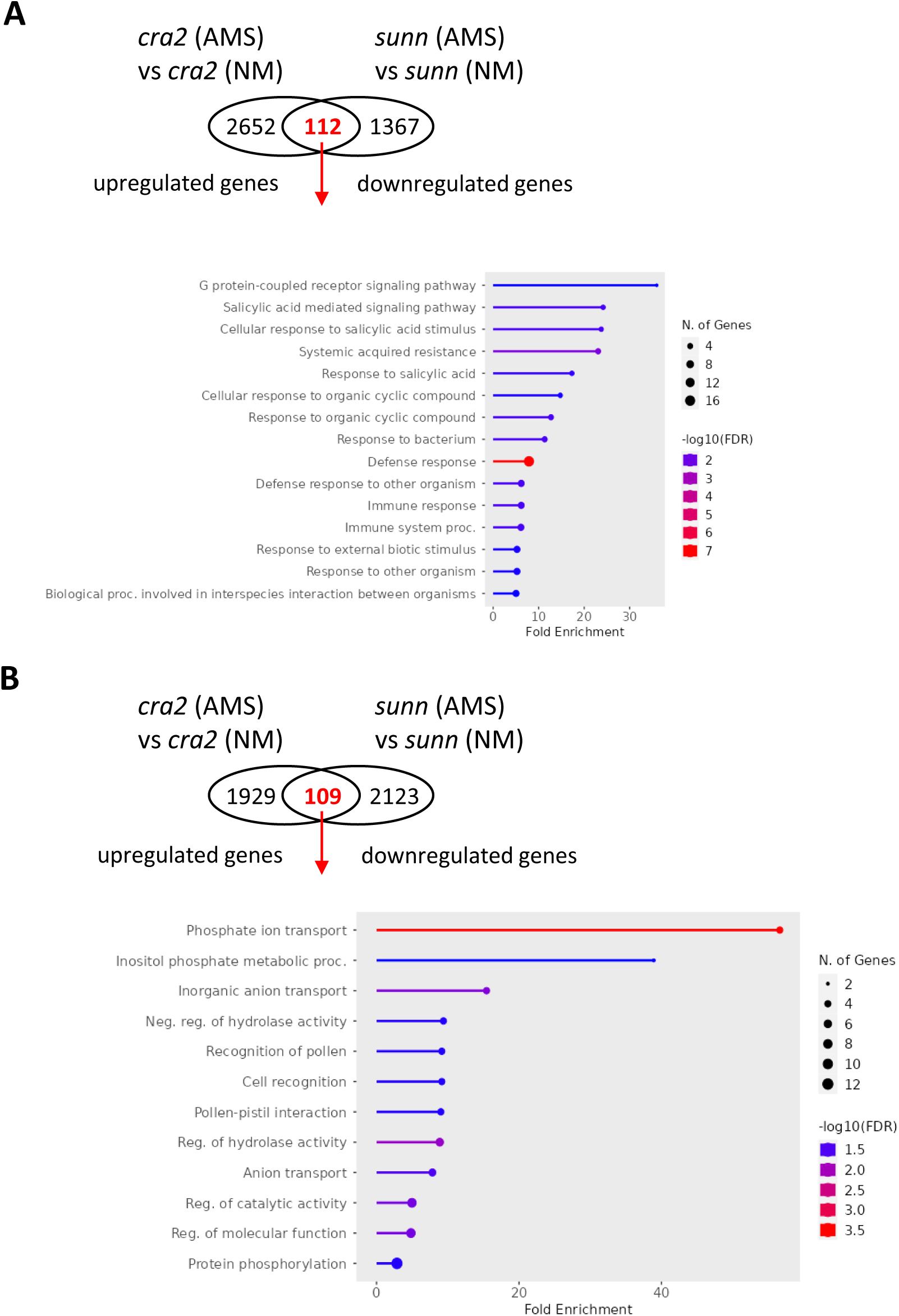
Genes antagonistically regulated in sunn and cra2 mutant mycorrhized roots (related to Figure 3) These comparisons were selected given the antagonistic role on the Arbuscular Mycorrhizal Symbiosis (AMS) of CRA2 (this study) and SUNN pathways^23,24^. Genes antagonistically regulated are listed in the Table S5. **A.** Venn diagram of genes upregulated in cra2 mutant roots grown with AMS fungi, based on the RNAseq dataset, and downregulated in sunn mutant roots grown in the same conditions, based on Karlo et al. (2020)^24^ data; and top 20 of enriched “Biological Process” Gene Ontology (GO) terms (hypergeometric distribution test, FDR < 0.05) observed in their intersection. **B.** Venn diagram of genes downregulated in cra2 mutant AMS roots, based on the RNAseq dataset, and upregulated in sunn mutant AMS roots, based on Karlo et al. (2020)^24^ data; and top 20 of enriched “Biological Process” GO terms (hypergeometric distribution test, FDR < 0.05) observed in their intersection.

**Figure S13.**
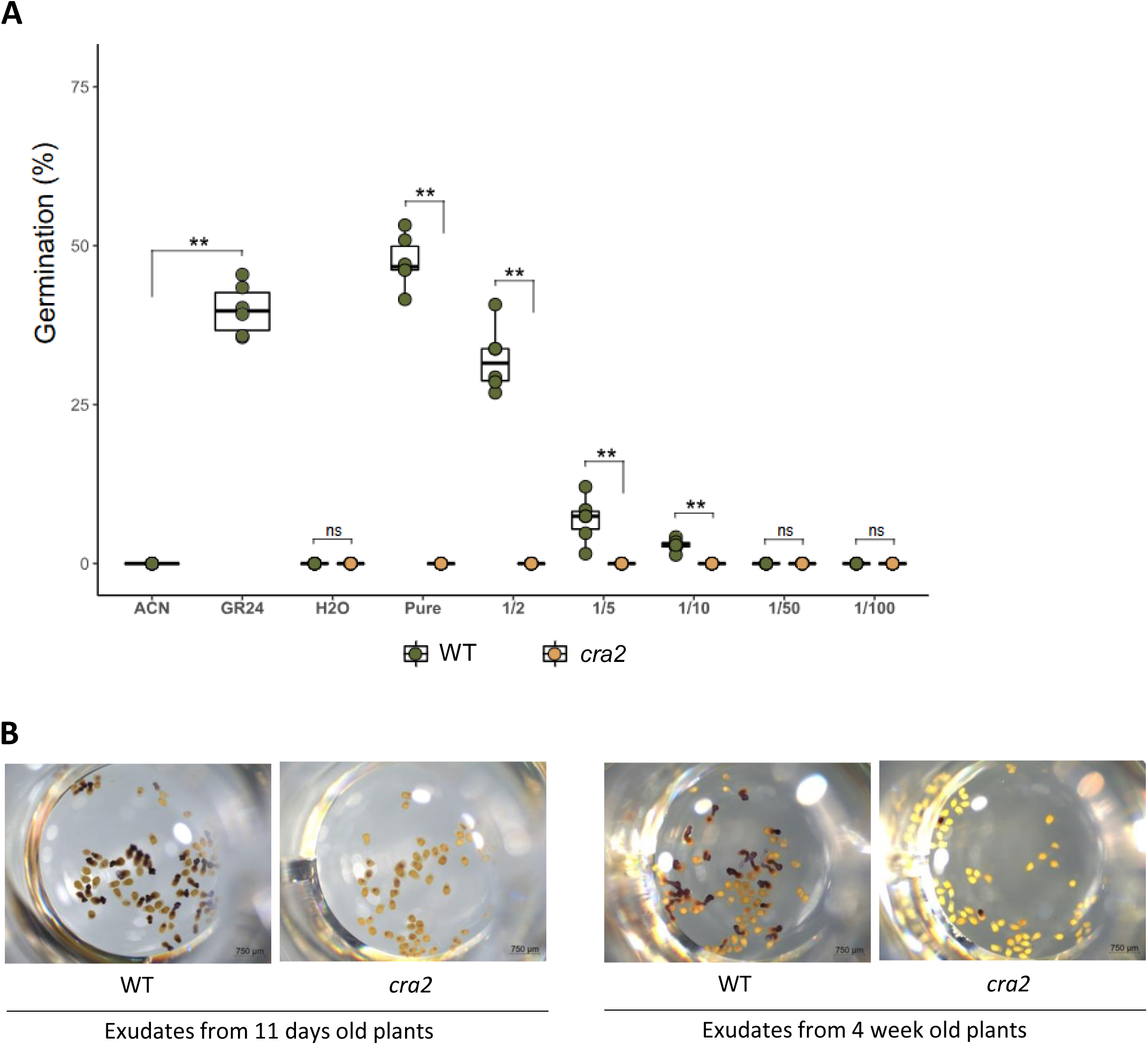
P. ramosa seeds response to root exudates of WT and cra2 mutant plants (related to Figure 4B) **A.** Eleven-days-old Wild-Type (WT) and cra2-11 mutant plants were grown under low Pi condition. A pool of five plants was transferred for 24h in sterile water to collect the root exudates and perform for P. ramosa seed germination assays (**A**). Boxes represent the middle 50% of the values, the median is represented by a horizontal line, and upper and lower quartiles by vertical lines. ACN, acetonitrile 0.001%; GR24, rac-GR24 10^−7^ M; pure, undiluted root exudates; 1/x : serial dilutions of pure root exudates with sterile water. A Mann-Whitney test was performed between WT and cra2 plants : **, P < 0.01. **B.** Representative pictures of Methylthiazolyldiphenyl-tetrazolium bromide (MTT) stained P. ramosa seeds treated with 11 days-old (left images) or 4-weeks-old (right images) M. truncatula Wild-Type (WT) and cra2-11 mutant plants root exudates.

**Table S1.** List of differentially expressed genes identified in the comparisons between cra2 mutant roots and Wild-Type roots under Non-Mycorrhizal or in Arbuscular Mycorrhizal Symbiotic conditions (related to Figure 3)

**Table S2.** List of genes associated to lateral root development that are differentially expressed in the cra2 mutant versus Wild-Type Non-Mycorrhizal roots (related to Figure 3)

**Table S3.** Fold change expression of MtSUNN, MtRAM2 and MtSTR2 mycorrhizal-related genes between cra2 mutant and Wild-Type roots in Non-Mycorrhizal or Arbuscular Mycorrhizal Symbiotic conditions (related to Figure 3)

**Table S4.** List of primers used in this study (related to Figure 1)

**Table S5.** List of Genes antagonistically regulated in sunn and cra2 mutant mycorrhized roots (related to Figure 3)

